# New microsatellite marker tools for genotype identification and analyses of genetic relationships in two ornamentals, the popular common lilac (*Syringa vulgaris*) and the invasive garden escapee Himalayan balsam (*Impatiens glandulifera*)

**DOI:** 10.1101/2020.03.03.974535

**Authors:** Helena Korpelainen, Maria Pietiläinen

**Affiliations:** Department of Agricultural Sciences, Viikki Plant Science Centre, P.O. Box 27 (Latokartanonkaari 5), FI-00014 University of Helsinki, Finland

**Keywords:** Genetic fingerprinting, Marker development by sequencing, *Impatiens*, *Syringa*, SSR markers

## Abstract

Nowadays, high-throughput sequencing technologies are widely available. Yet, it is practical to have an access to simpler and cheaper, yet effective low-throughput analyses as well. For that purpose, species-specific microsatellites, also called simple sequence repeats (SSR), are valuable, multi-purpose types of markers. In the present study, we introduce new sets of SSR markers for two ornamental plant species, the popular common lilac (*Syringa vulgaris* L.) (16 markers) and the invasive garden escapee Himalayan balsam (*Impatiens glandulifera* Royle) (259 markers). The markers were developed as a by-product of a genotyping-by-sequencing project producing a large amount of DNA sequence data. Both the frequency of SSRs and the success rate for marker development were considerably greater in *I. glandulifera* when compared to *S. vulgaris*. The new markers will contribute to the characterization of germplasm and to other types of genetic analyses on these two species.

## 1. Introduction

DNA markers are a powerful tool for assessing genetic diversity, differences and characteristics within a species, and for the identification and fingerprinting of genotypes. Practical applications range from marker-assisted breeding to environmental conservation, parentage analyses and forensic investigations. Despite a wealth of available DNA information, there are numerous species with lacking or a narrow range of adequate genetic markers. Nowadays, high-throughput sequencing can be applied to most everything, but it is practical to have an access to simpler and cheaper, yet effective low-throughput analyses as well. For that purpose, species-specific microsatellites, also called simple sequence repeats (SSR) and composed of short tandem repeats, are valuable, multi-purpose types of markers. In the present study, we will introduce new sets of SSR markers for two ornamental plant species, the popular common lilac (*Syringa vulgaris* L.) and the invasive garden escapee Himalayan balsam (*Impatiens glandulifera* Royle). At the present, there are only 9 and 11 species-specific SSR markers available for *S. vulgaris* (JuntheikkiPalovaara et al., 2013) and for *I. glandulifera* (Provan et al., 2007; Walker et al., 2009), respectively.

*S. vulgaris* (Oleaceae) is a popular ornamental shrub or a small tree, native to the Balkans (Lack, 2001). It was first cultivated in Central Europe in the sixteenth century, but it quickly found its way to gardens in Western Europe and later to gardens in temperate regions all over the world (Lack, 2001). The precise identity of *S. vulgaris* accessions may be unknown and difficult to solve. Therefore, breeders, producers, retailers and consumers will benefit from the development of efficient molecular markers for a precise identification and characterization of cultivars. While *S. vulgaris* is a popular ornamental, *I. glandulifera* (Balsaminaceae) is a tall annual plant, originating from the Himalayas and presently considered an invasive plant that grows rapidly and spreads effectively. It was introduced to Europe in 1839 as a garden ornamental (Beerling and Perrins, 1993). Thereafter, *I. glandulifera* has spread widely throughout Europe (Beerling and Perrins, 1993), and it occurs also in North America and New Zealand as an invasive plant (Weber, 2003). Despite its beautiful flowers, the main interest is not to cultivate but to control and eradicate *I. glandulifera* populations. Its success may be due to its previous popularity as an ornamental garden plant, its rapid growth rate, good ability to survive heavy frost and high seed production (Perrins et al., 1993). Population genetic studies utilizing DNA markers may provide better understanding about the drivers of dynamic invasion processes and about the evolutionary opportunities that invasive plants have acquired. Such knowledge combined with life history studies can aid management practices.

## 2. Materials and methods

In all, 85 *S. vulgaris* genotypes (additional 9 samples failed in the analyses) including international reference samples, historical samples from Finland, Sweden and France, and unidentified cultivars from Finland were exposed to genotyping-by-sequencing (GBS) to produce DNA sequence data and to perform simultaneously analyses based on SNP and SilicoDArT markers to allow a genome-wide study on genetic diversity and differentiation (Korpelainen et al., in preparation). SNP markers are nucleotide polymorphisms present in the tag sequences, while SilicoDArT markers represent presence/absence variation (PAV) in the tag sequences. Comparably, 84 *I. glandulifera* samples (additional 10 samples failed in the analyses) originating from Finland, England, Canada, India and Pakistan were subjected to similar analyses as *S. vulgaris* (Korpelainen and Pietiläinen, in preparation). At the same time, the sequence data obtained from these two species allowed the discovery of microsatellite repeat regions and primer development to generate sets of SSR (microsatellite) markers for further use in simpler and cheaper low-throughput analyses on *Syringa* and *Impatiens*.

Prior to SNP and SilicoDArT analyses, genomic DNA was extracted from leaf tissue using the CTAB protocol of Doyle and Doyle (1990) or a commercial kit (E.Z.N.A.™ Plant DNA Mini Kit Spin Protocol (Omega Bio-Tek). The quality and quantity of extracted DNA were quantified with a spectrophotometer and further confirmed on 0.8% agarose gels. DNA concentrations were adjusted at 50 ng μl^-1^. DNA samples of *S. vulgaris* and *I. glandulifera* were sent to Diversity Arrays Technology Pty Ltd. (Canberra, Australia; http://www.diversityarrays.com) for high-throughput sequencing utilizing the DArTseq system and for marker discovery (Kilian et al., 2012).

The sequence data consisted of 69 bp long pieces of data. All available sequences were used to explore for SSR repeats using MSATFINDER version 2.0.9. (Thurston and Field, 2005) combined with visual observations with the following criteria: mononucleotide repeats with at least eight copies and other types of repeats with a minimum of five copies. All SSRs fulfilling the criteria were recorded. The Primer3 software (Rozen and Skaletsky, 2000) was used to design primer pairs for the potential amplification of the discovered SSRs when possible (SSR located at a relatively middle position of the tag sequence containing a SSR, and the primers being 18-27 bp in length and with the GC content 40-60%

## 3. Results and discussion

The total length of the sequenced area obtained in SNP and DArTseq analyses equalled 1 035 552 bp and 740 508 bp, respectively, in S. *vulgaris*, and 1 447 827 bp and 2 044 125 bp, respectively, in *I. glandulifera*. The numbers of SSR regions found in *S. vulgaris* based on SNP and DArTseq analyses equaled 111 (repeat motifs: 63.1% mononucletide, 31.5% dinucleotide and 5.4% compound) and 116 (repeat motifs: 82.8% mononucletide, 13.8% dinucleotide and 3.4% compound), respectively (Table 1). In *I. glandulifera*, the numbers of SSRs found based on SNP and DArTseq analyses equaled 477 (repeat motifs: 76.9% mononucletide, 11.9% dinucleotide, 8.4% trinuleotides, 0.2% tetranucleotides, 0.6% pentanucleoties and 1.9% compound) and 709 (repeat motifs: 73.9% mononucletide, 11.8% dinucleotide, 11.6% trinucleotide, 0.8% tetranucleotide, 0.4% pentanucleotide, 0.1% hexanucleotide and 1.3% compound), respectively (Table 2). Thus, the great majority of all found SSRs were composed of mononucleotide repeats, followed by dinucleotide repeats, while repeat motifs longer than three nucleotides were very rare. The frequency of SSRs relative to the total sequence length was considerably higher in *I. glandulifera* than in *S. vulgaris*, about triple and double in SNP and DArTseq analyses, respectively. The relative frequency of SSRs was slightly higher in SNP-based analyses when compared to DArTseq-based analyses.

**Table 1.** Numbers and percentages of different microsatellite repeats found in 1 035 552 bp and 740 508 bp of DNA sequences obtained from SNP and SilicoDArT marker analyses, respectively, in *Syringa vulgaris*.

| SNP analysis |  |  | SilicoDArT analysis |  |  |
| --- | --- | --- | --- | --- | --- |
| Repeat motif | Number | s % repeats | Repeat motif | Numbers | % repeats |
| A<sub>8</sub>/T<sub>8</sub> | 54 | 48.6 | A<sub>8</sub>/T<sub>8</sub> | 52 | 44.8 |
| A<sub>9</sub>/T<sub>9</sub> | 5 | 4.5 | A<sub>9</sub>/T<sub>9</sub> | 22 | 19.0 |
| A<sub>10</sub>/T<sub>10</sub> | 4 | 3.6 | A<sub>10</sub>/T<sub>10</sub> | 14 | 12.1 |
| A<sub>11</sub>/T<sub>11</sub> | 2 | 1.8 | A<sub>11</sub>/T<sub>11</sub> | 5 | 4.3 |
| A<sub>13</sub> | 1 | 0.9 | A<sub>12</sub>/T<sub>12</sub> | 2 | 1.7 |
| A<sub>15</sub> | 1 | 0.9 | G<sub>9</sub> | 1 | 0.9 |
| T<sub>16</sub> | 1 | 0.9 | [A/G]<sub>22</sub> | 1 | 0.9 |
| T<sub>17</sub> | 1 | 0.9 | [AC]<sub>6</sub> | 2 | 1.7 |
| T<sub>18</sub> | 1 | 0.9 | [AC]<sub>10</sub> | 1 | 0.9 |
| [AG]<sub>5</sub> | 1 | 0.9 | [AG]<sub>6</sub> | 1 | 0.9 |
| [AT]<sub>5</sub> | 1 | 0.9 | [CA]<sub>10</sub> | 1 | 0.9 |
| [AT]<sub>6</sub> | 1 | 0.9 | [CG]<sub>5</sub> | 1 | 0.9 |
| [AC]<sub>5</sub> | 2 | 1.8 | [CT]<sub>8</sub> | 1 | 0.9 |
| [AG]<sub>10</sub> | 1 | 0.9 | [GA]<sub>8</sub> | 1 | 0.9 |
| [AG]<sub>15</sub> | 1 | 0.9 | [GA]<sub>13</sub> | 1 | 0.9 |
| [CA]<sub>5</sub>/[TG]<sub>5</sub> | 7 | 6.3 | [GT]<sub>5</sub> | 1 | 0.9 |
| [CA]<sub>6</sub> | 2 | 1.8 | [GT]<sub>7</sub> | 1 | 0.9 |
| [GA]<sub>5</sub> | 4 | 3.6 | [TC]<sub>5</sub> | 3 | 2.6 |
| [GA]<sub>6</sub> | 3 | 2.7 | [TC]<sub>6</sub> | 1 | 0.9 |
| [GA]<sub>8</sub> | 1 | 0.9 | [TG]<sub>5</sub> | 1 | 0.9 |
| [GA]<sub>12</sub> | 1 | 0.9 | [GT]<sub>5</sub>[GA]<sub>4</sub> | 1 | 0.9 |
| [GA]<sub>14</sub> | 1 | 0.9 | T<sub>4</sub>CT<sub>8</sub> | 1 | 0.9 |
| [GA]<sub>30</sub> | 1 | 0.9 | G<sub>9</sub>AG<sub>5</sub> | 1 | 0.9 |
| [TC]<sub>5</sub> | 7 | 6.3 |  |  |  |
| [TC]<sub>11</sub> | 1 | 0.9 |  |  |  |
| T<sub>8</sub>CT<sub>4</sub> | 1 | 0.9 |  |  |  |
| A<sub>8</sub>TA<sub>7</sub> | 1 | 0.9 |  |  |  |
| A<sub>5</sub>N<sub>2</sub>A<sub>4</sub>NA<sub>8</sub> | 1 | 0.9 |  |  |  |
| A<sub>2</sub>TA<sub>8</sub>TA<sub>3</sub> | 1 | 0.9 |  |  |  |
| A<sub>6</sub>TA<sub>8</sub> | 1 | 0.9 |  |  |  |
| T<sub>6</sub>N<sub>6</sub>T<sub>8</sub> | 1 | 0.9 |  |  |  |
| Total | 111 |  | Total | 116 |  |

**Table 2.**
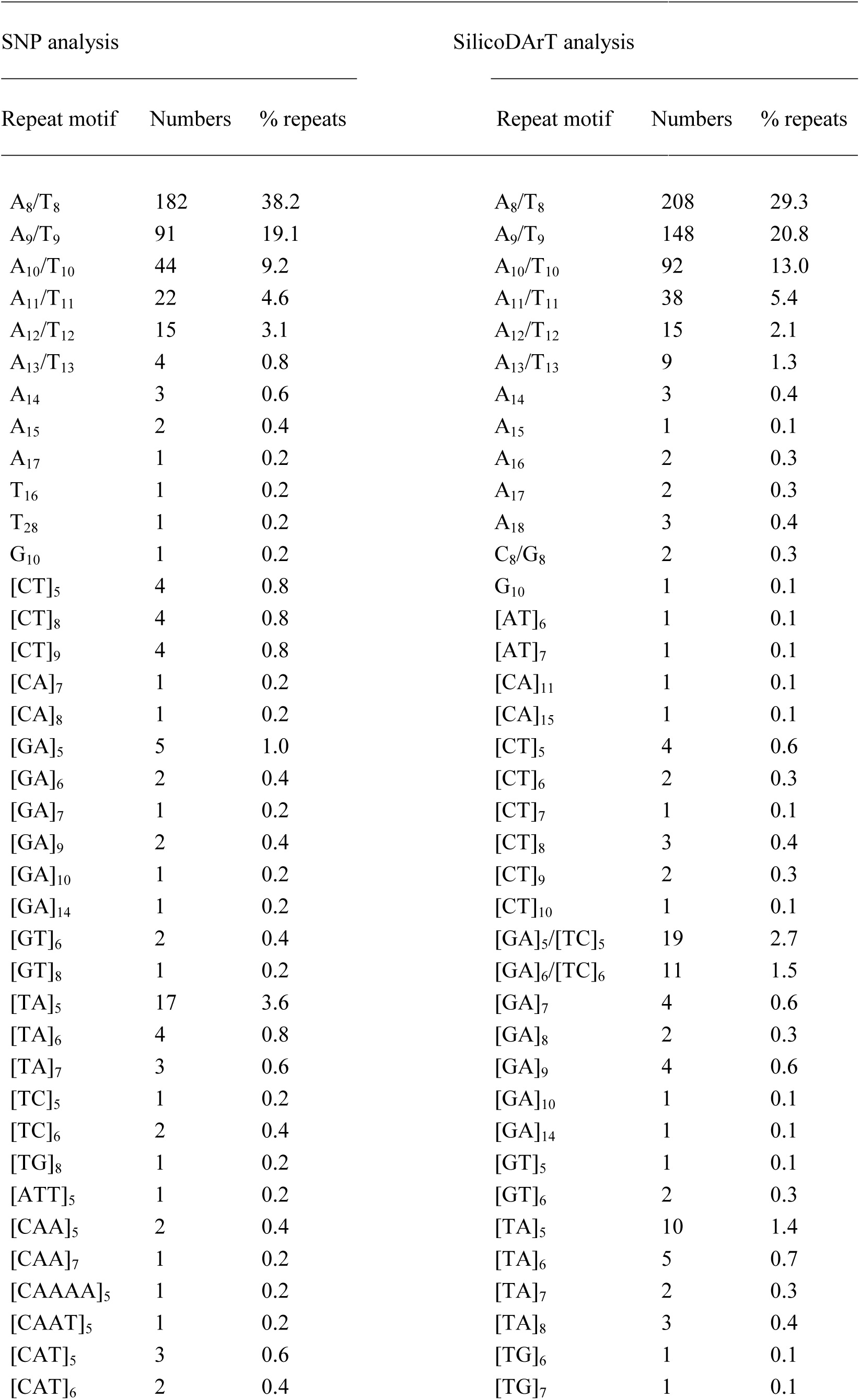
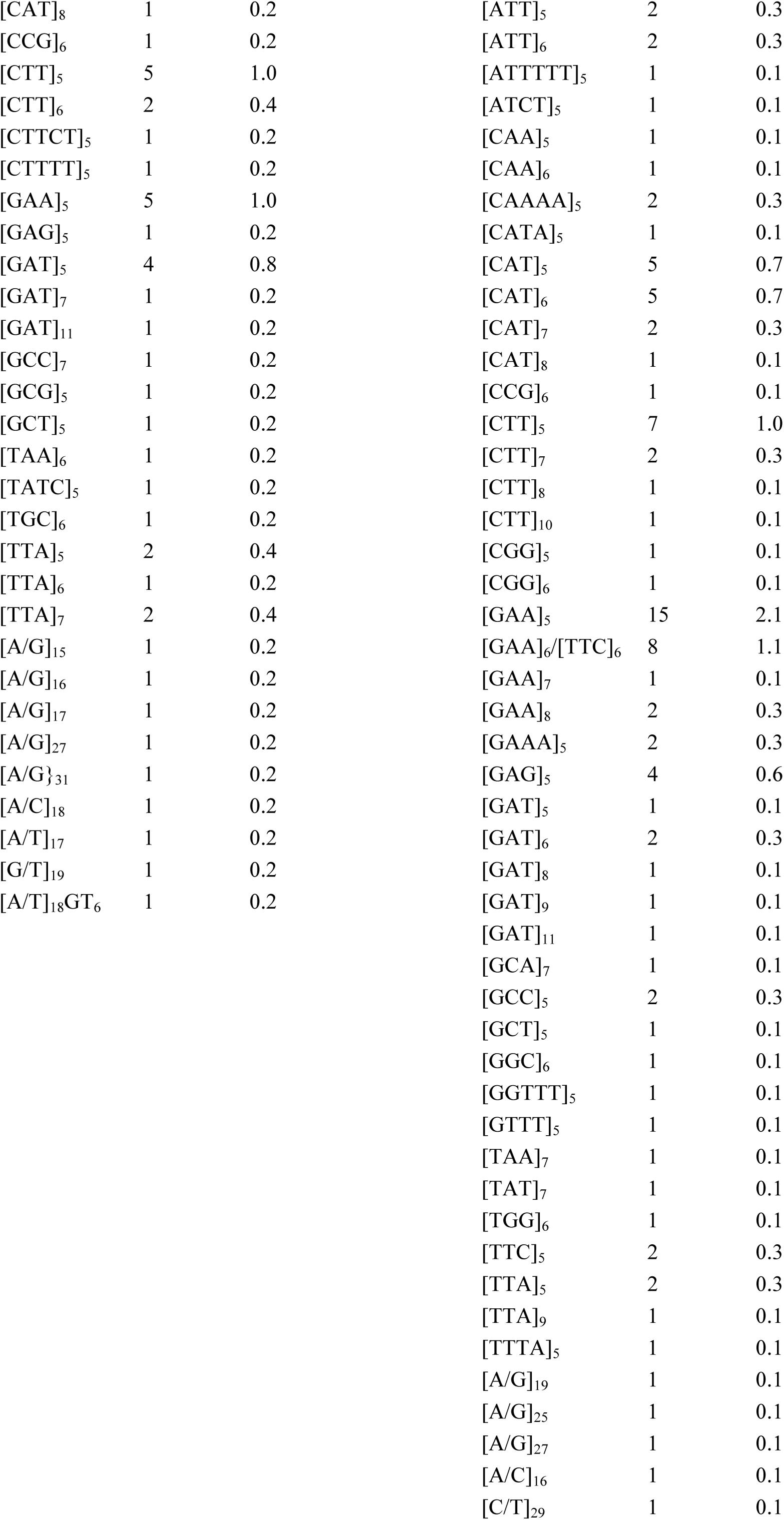
Numbers and percentages of different microsatellite repeats found in 1 447 827 bp and 2 044 125 bp of DNA sequences obtained from SNP and SilicoDArT analyses, respectively, in *Impatiens glandulifera.*

We succeeded to design adequate primer pairs for only a portion of the discovered SSRs: in *I. glandulifera*, for 23.5% (112 primer pairs) and 20.7% (147 primer pairs) of SNP- and DArTseqbased SSRs, respectively, while in *S. vulgaris*, the percentages were much lower, only 14.4% (16 primer pairs) and 0% of SSRs, respectively (Tables S1-S2). Thus, both the frequency of SSRs and the success rate for marker development were considerably greater in *I. glandulifera* when compared to *S. vulgaris*.

Previous DNA-based studies on the genus *Syringa* have involved the use of RAPD markers (Chen et al., 1999; Kochneva et al., 2004; Melnikova et al., 2009), ISSR markers (Rzepka-Plevneš et al., 2006; Yang et al., 2013), AFLP markers (Ming and Gu, 2006), sequencing of nuclear and chloroplast regions (Smolik et al., 2010; Lendvay et al., 2016), and recently the sequencing of the whole chloroplast genome of *S. pinnatifolia* (Zhang et al., 2019). More effective and reliable SSR markers for cultivar identification in *S. vulgaris* have been developed by Juntheikki-Palovaara et al. (2013, nine microsatellites). In addition, Lendvay et al. (2013) have developed five microsatellites for *S. josikaea*, and De La Rosa et al. (2002) have reported that some markers developed for olive (*Olea europaea* L.) amplified also in the genus *Syringa*. Although these markers appeared valuable for detecting differentiation among cultivars, the availability of powerful markers has been limited to allow high-precision cultivar identification. The 16 SSRs introduced in the present study are a useful addition to the 9 species-specific markers previously available for *S. vulgaris* (JuntheikkiPalovaara et al., 2013). They will contribute to the characterization of germplasm and to other types of genetic analyses in *S. vulgaris* and possibly in other species of the genus *Syringa*.

Considerable research attention has been paid on *I. glandulifera*, which is a major invasive plant that spreads effectively. Previously, Provan et al. (2007) have developed eight SSR markers for *I. glandulifera*, and Walker et al. (2009) additional three markers. These markers have been used in population genetic studies by Love et al. (2013), Hagenblad et al (2015), Nagy and Korpelainen (2015), and Helsen et al. (2019). *I. glandulifera* has also been the target of RAPD analyses (Zybartaite et al., 2011; Kupcinskiene et al., 2015) and ISSR analyses (Kupcinskiene et al., 2015). The new SSRs (a total of 259 markers) are a huge increase in the selection of markers that could be used in different types of genetic studies on *I. glandulifera*. However, testing the true efficiency of each marker in genetic analyses is beyond the present short communication paper.

## Acknowledgements

The project was supported by the Nikolai and Ljudmila Borisoff Foundation. We would like to thank Juulia Kotkavuori and Leena Lindén for advice and help in sampling materials.

## Supplementary data

Supplementary material related to this article: Table S1 and Table S2.

