## Supplemental Table 1 for "New microsatellite marker tools for genotype identification and analyses of genetic relationships in two ornamentals, the popular common lilac (*Syringa vulgaris*) and the invasive garden escapee Himalayan balsam (*Impatiens glandulifera*)"

**Table S1 Supplementary material**

Characteristics and primer sequences of SSR markers developed for *Syringa vulgaris*. T_a_, annealing temperature. The markers were developed based on sequence information obtained in high-throughput SNP analyses.

________________________________________________________________________________________________

Locus	Repeat motif	Primer sequences (5’-3’)	T_a_	Product size (bp)	Accession number

__________________________________________________________________________________________________________

| SyrA1 | [GA]_5_ | F: AGGGACCTGTGAGTAATTGATTGA | 59 | 53 | MK520881 |
| --- | --- | --- | --- | --- | --- |
|  |  | R: TTGGGAGGAACAAGTGGTGG |  |  |  |
| SyrA2 | [AG]_5_ | F: TGCAGTTGATGGACCATACAGA | 59 | 59 | MK520882 |
|  |  | R:TGTCCCTCAAATCAGTGTAGATAGAC |  |  |  |
| SyrA3 | [GA]_5_ | F: TGCAGTAACCTCCATATCAGTAACT | 59 | 67 | MK520883 |
|  |  | R: ACCAAAGTTCTACCCTACAGAGC |  |  |  |
| SyrA4 | [GA]_14_ | F: TGCAGGTTTGACAGAGAGAGAG | 58 | 60 | MK520884 |
|  |  | R: TTGTCCAGGGGCCATTTCTC |  |  |  |
| SyrA5 | [GA]_8_ | F: TGCAGATTCGGTTGAGGGAG | 59 | 65 | MK520885 |
|  |  | R: CTCTCACTCCCGACATCTCTC |  |  |  |
| SyrA6 | [TC]_5_ | F: TGCAGTCAGAAAATCTCCTCCA | 58 | 60 | MK520886 |
|  |  | R: ACTCCCTGTTATTCAAGAAATGTAAA |  |  |  |
| SyrA7 | A_8_ | F: GCAGCTTTGGGTTATTGCAAC | 58 | 57 | MK520887 |
|  |  | R: TGCATGTAGGGAATCTGAAGGA |  |  |  |
| SyrA8 | A_8_ | F: GGGGTTCAGATGGGATTTTACA | 58 | 64 | MK520888 |
|  |  | R: TCATTTCAACATTGCATCACCTT |  |  |  |
| SyrA9 | A_8_ | F: CCCGTAGAACTGTGTTGTGC | 59 | 56 | MK520889 |
|  |  | R: CCAAAGTACAGAATAAGTCCCAACC |  |  |  |
| SyrA10 | A_9_ | F: TGCAGTTTCCTTACATGTATGCC | 58 | 59 | MK520890 |
|  |  | R: TGGTCAATTCTATCCTCGTCATT |  |  |  |
| SyrA11 | A_8_ | F: AGCCAAAGGTTGAAATGAGGA | 58 | 63 | MK520891 |
|  |  | R: TGTGTGGACTCGGGACCTAT |  |  |  |
| SyrA12 | A_8_ | F: CAGAGACTGGTAGGGAGTAAGT | 58 | 55 | MK520892 |
|  |  | R: TGACACCCCTATAGCACAATTT |  |  |  |
| SyrA13 | T_6_N_6_T_8_ | F: GCAGCAGTCCACCCATAGTT | 60 | 66 | MK520893 |
|  |  | R: CGCAAGGGAAAACACCAGTG |  |  |  |
| SyrA14 | T_9_ | F: AGTTTCAGTCTGGGTTGGCA | 59 | 63 | MK520894 |
|  |  | R: GTCATTGGTAATATGCCTGGTGG |  |  |  |
| SyrA15 | T_8_ | F: TGCAGTTCCGATCTCTCAGG | 58 | 62 | MK520895 |
|  |  | R: ACATTTACTAATCTAGGACTAAAGGGG |  |  |  |
| SyrA16 | [GA]_6_ | F: GCAAATGTGAGCGAGCGAG | 59 | 55 | MK520896 |
|  |  | R: CACACTCCTTTCTCTCTTTCATCG |  |  |  |

_________________________________________________________________________________________________________
