## Supplemental Table 2 for "New microsatellite marker tools for genotype identification and analyses of genetic relationships in two ornamentals, the popular common lilac (*Syringa vulgaris*) and the invasive garden escapee Himalayan balsam (*Impatiens glandulifera*)"

**Table S2 Supplementary material**

Characteristics and primer sequences of SSR markers developed for *Impatiens glandulifera*. Ta, annealing temperature. ImpA and ImpB markers were developed based on sequence information obtained in high-throughput SNP and SilicoDArT analyses, respectively.

__________________________________________________________________________________________________________

Locus	Repeat motif	Primer sequences (5’-3’)	Ta	Product size (bp)	Accession number

__________________________________________________________________________________________________________

ImpA1	A_8_	F: TCTCCTATATTCAGTTATCTTGA	53	55	MK688998

R: ACCCCAAATTTCCTTGAAGT

ImpA2	[CTT]_5_	F: TGCAGCACCTCAACCATTTC	59	57	MK688999

R: GTCATGAACTCCCAATGAAGAAGA

ImpA3	[TA]_5_	F: ACTGTACTCCTCTCTTTCTCT	55	63	MK689000

R: GCCTATGTATCCGATTATAAGCT

ImpA4	T_8_	F: TGCAGTTATCCTTTTACCTAAA	55	68	MK689001

R: GGAACTAGAAAGACAGAGAGGG

ImpA5	A_8_	F: TGCAGGGATGAGGGTTTATTG	58	69	MK689002

R: GGTTCTCTCAAATAATTCAACCGA

ImpA11	T_9_	F: CAGACATATACTGGAACCTT	52	65	MK689003

R: TCTGTAAAATGGTGTTGTACA

ImpA12	T_9_	F: TGCAGCCATAACTACAAAGCT	58	66	MK689004

R: CTGTAATTTGGTCCGTCTGCA

ImpA13	[TA]_5_	F: TGCAGCGCCGACCTAGAG	60	57	MK689005

R: TCCCTTGTCCATTTGTTTACCC

ImpA14	A_10_	F: TGCAGGGATGAGGGTTTATTG	56	69	MK689006

R: GTTCTCTCAAATAATTCAACCGA

ImpA15	A_10_	F: TGCAGGGATGAGGGTTTATTG	56	69	MK689007

R: GTTCTCTCAAATAATTCAACCGA

ImpA16	[CT]_5_	F: TGCAGCCGACTATATTCTTCA	56	69	MK689008

R: CATGTAACAAGTGGAGGATGA

ImpA18	A	F: GAAAGAATCTTCAGCATTTTGA	54	55	MK689009

R: GTTTGAAACTCATCCACATGT

ImpA19	A_8_	F: TGCAGACACAATCATCGAT	55	66	MK689010

R: GCTCTACCAATGACTGTGCT

ImpA20	T_8_	F: TGCAGACACCCTCTCTTATT	56	64	MK689011

R: AGCCTTTACAATCTTCAACTTTCA

ImpA21	[GA]_14_	F: TGCAGAGGTTACAGTAGAGAGAG	57	68	MK689012

R: GCGACAATTTCAAGCTCTCT

ImpA22	[CT]_8_	F: AATTTACCTTCAAACAAGACTG	54	60	MK689013

R: CACAGGAGAGACTAGAGAGAGA

ImpA23	[TA]_6_	F: TGCAGATGAATGATGAATGATGA	55	60	MK689014

R: TGCATGCACCAAAATATCTT

ImpA24	A_8_	F: TGCATATCATCATAGGGAAGCA	57	58	MK689015

R: AATCCACCATCTTCTGTCGT

ImpA25	[GA]_9_	F: GGTCACGCCGCCTGAGAGAG	62	50	MK689016

R: GCGCCCCTCTCTCTCTCTTT

ImpA26	T_8_	F: GCAGTAAAACCAGAATATCCA	52	68	MK689017

R: GAATCAGAGACCAAGAAAA

ImpA27	A_8_	F: TGCAGCAATCATTCACATG	54	67	MK689018

R: ACATCTGTCATTTCCTCTTCT

ImpA28	A_10_	F: TGCAGCAATCCAAAAGCATA	54	69	MK689019

R: CCGATCTGTAATTTCTCCTT

ImpA29	[GA]_5_	F: GCAGCCACAGAGAGAGATTTG	59	63	MK689020

R: CCTTCTTCTTCAATTGAACCCCA

ImpA30	T_9_	F: GTCGCCTGGAAGCTTGATAC	57	60	MK689021

R: TCAAATTCCATTTACCAACTCCA

ImpA31	[CAAAA]_5_	F: TGCAGCTCATTGAAACATAACA	54	68	MK689022

R: TCTTCTTCTGTATTTTCTCTGT

ImpA32	[TA]_5_	F: CAGCTTCGTTCCCACCAAAA	56	67	MK689023

R: CGATCTGTAATGATGATAAATGCA

ImpA33	[CAA]_5_	F: GCAGGAGATCACAGCAACAA	56	68	MK689024

R: TGCCAGATCTCATCATCATT

ImpA34	[GAT]_7_	F: GTAGTAGTACTCAGACTTGGAAG	56	55	MK689025

R: ACTGCCACCCATCATCATCA

ImpA35	A_9_	F: TGCCAAATTCTTCATTCTGC	52	56	MK689026

R: GAAGATATTCCTAATACAACATC

ImpA36	A_8_	F: GTAACCTTTGCCTGTGAGTA	52	54	MK689027

R: AACTACAAATGCTAAACCAT

ImpA37	T_8_	F: GTTCGAATCCTACTTTGGG	52	60	MK689028

R: AAAAGGTCATTTGGGAAAA

ImpA38	T_8_	F: TGCAGGTTCTCAATTTATTTCTCT	56	62	MK689029

R: GCAAAACAGCTAATGAATTCTCT

ImpA39	A_8_	F: TGCTTCCATCCAAAAGAAAA	52	59	MK689030

R: GCAGACAAAACAACATAAAT

ImpA40	A_9_	F: TGCAGTAACAGAAAATGAAA	53	64	MK689031

R: CTCTCTAGGCAAATTGATCAGA

ImpA41	[G/A]_17_	F: AATGGAGGTAAGACGTGGGG	57	63	MK689032

R: CCTTCTTCTTTAGCTCAACTTCT

ImpA42	[C/A]_18_	F: CAGTACTGCATGCACAAGA	53	55	MK689033

R: TTATAGTACATTATAGCTGTGTG

ImpA43	T_8_	F: TGCAGTGAAGTTTCTTCCAC	55	58	MK689034

R: AAAAGCATCTTTGGAAGTCC

ImpA44	A_9_	F: TGCAGACCTGTTTGTTTGTGT	59	67	MK689035

R: ATGGCATGGGATGGGAAGAC

ImpA45	T_9_	F: TGCAGCCCTACTATTTGAAAGC	56	65	MK689036

R: CTGTAAGTTGTAGGTCCAAAAT

ImpA46	[CT]_8_	F: GCAGGAAGAAGAAATGTGGC	54	68	MK689037

R: TGTAAATTGAGACAATACTAGGA

ImpA47	T_10_	F: GCAGTTGGTTGTGATCTTCA	54	57	MK689038

R: TCTAGCTGTTATCCAAATAGAT

ImpA48	A_8_	F: TTCTGTTCTGAAGGCCGAGG	57	63	MK689039

R: CGCCTGTTCTCATCTATCAA

ImpA49	[GAG]_5_	F: TGCAGGAAATTGTTGTAT	51	66	MK689040

R: TAACAACCCCTTCTCTTTCT

ImpA50	[CAT]_8_	F: TGCAGCAAGAAGATCCATCA	57	69	MK689041

R: AGGTAAGTCTCTTTCATACTGCT

ImpA57	A_8_	F: GCAGTCATGTCATCTAATCAGGA	55	68	MK689042

R: CTTCCGATCTGTAAACACAA

ImpA60	T_8_	F: ACAGGGGAGATTCAATGAAA	54	57	MK689043

R: CTCTTCCGATCTGTAATCCA

ImpA65	T_10_	F: TGCAGAAACATTCAAAGCCT	56	60	MK689044

R: GGCATCTCTCAAGGTTGTGT

ImpA66	T_8_	F: TGCAGAAACCAAATCAATTT	53	66	MK689045

R: CTGTAATAGCCAGTGTAATGCA

ImpA67	T_10_	F: TGCAGAAAGGTAAAGATGTATG	52	67	MK689046

R: TCCAAAACATTTAGAGTGAA

ImpA68	[TTA]_5_	F: ACTGGTTGGTTCATTTAGGT	52	63	MK689047

R: AAATTACAACACTAAAGGCT

ImpA70	T_10_	F: CAGACATATACTGGAACCTT	52	67	MK689048

R: CGATCTGTAAAATGGTGTTGT

ImpA71	[TA]_5_	F: CAGCCAACTTTCTCCTGAA	52	59	MK689049

R: CGATTTGTGTTGTTTCATTA

ImpA72	[A/G]_16_	F: GCAGATCAAAACATTTTACA	50	57	MK689050

R: TTACCTACGAAGATTCTTCT

ImpA73	[A/G]_15_	F: TGCAGATGATAAGTATGAAGA	51	61	MK689051

R: AAAGAACTCAACAACTTTCT

ImpA74	[G/A]_27_	F: TGCAGATTACGACGTTATTT	54	65	MK689052

R: CGGTCATCGTCGTAATCTTT

ImpA75	[TA]_5_	F: CAAAATGGGCTGTACAAATA	51	56	MK689053

R: CCGATCTGTAACATAATGAA

ImpA76	[A/T]_18_GT_6_	F: CAGATGCTCTCGGTTAGA	52	63	MK689054

R: GATGGATGATCAAACACAAA

ImpA77	T_8_	F: CTGCTGCTTGACAAACTGAA	57	54	MK689055

R: ACCATTTTACATGTAGATCCGACA

ImpA78	[CAT]_6_	F: TGGCTACCACCTTATCATCA	56	54	MK689056

R: GATTATGAATTTCGTCCTTGAATGA

ImpA79	T_8_	F: TGCAGCTCCTTGATCTCAA	55	66	MK689057

R: TGTCATCATTTGCATTTTAGAAG

ImpA80	[CTT]_6_	F: GCATCTTCAACAGCTTCTTCT	53	65	MK689058

R: AGAAGAAGAAGATGATCGTC

ImpA81	[CTT]_6_	F: CTTCTTCAACAGCTTCTTCTTCT	54	54	MK742816

R: GATGATCGTCAGCCAAAG

ImpA82	[CAT]_5_	F: CAGCTTTCCCCTGTCTCATT	57	63	MK742817

R: TGTAGGAGGGTAGAGAAGGACA

ImpA83	[GAA]_5_	F: GGAGGTTGCAGGACTTATTC	52	55	MK742818

R: GCCAGAATAAGCAACTTC

ImpA84	A_8_	F: TGCAGTCAGATCAATTGGAA	55	62	MK742819

R: ACCAATATCATTCAATCCTCCGT

ImpA85	A_8_	F: GGAAACCTGTCAACAAGATCA	54	54	MK742820

R: TCTTTGTTTTCGTGTTTGAA

ImpA86	T_9_	F: AACTTGACCCCAAAGTAGAA	52	58	MK742821

R: TGTTCTCACTAAATCTGAAA

ImpA87	[G/A]_31_	F: TGCAGAAAAGAAGCGGGC	58	62	MK742822

R: CCCCTTCAAATCATCGACCAG

ImpA88	A_10_	F: TGCAGACCTGTTTGTTTGTGT	59	68	MK742822

R: ATGGCATGGGATGGGAAGAC

ImpA89	T_8_	F: TGAAACAGAGAGAAAGCGTA	52	57	MK742824

R: CATGCTAGTTTCCTACAATAA

ImpA90	T_9_	F: TGCAGTCTGTCGGTTTATAAGA	57	69	MK742825

R: ACTCCATTTGTCATCCCAAGT

ImpA91	[TA]_6_	F: TGCGCTGAATCCTTCTAAAG	53	62	MK742826

R: GATCTGTAATCGAAATGGTT

ImpA92	[TC]_5_	F: AGTTCCACATAGTAATCCGATTATT 56	58	MK742827

R: AGAAATCGAGCGGTGAGAGA

ImpA93	T_11_	F: GTTGGCTGTGATCTTCAATGGA	56	59	MK742828

R: TCTAGTGCTGTTATCCAAATAGA

ImpA94	[A/T]_17_	F: TGCAGCCAAAATAGATTATAGTTGA 57	69	MK742829

R: GTAGTTCGACCCCTACCTGG

ImpA100	T_9_	F: TGCAGTAGTATAAGACAGTAATGA 55	58	MK742830

R: CTGTAAGTCTCGTTTGTAAGCA

ImpA102	A_8_	F: TGCAGTTCTAGAAACTTTAGA	54	69	MK742831

R. TCTTCCGATCTGTAATGAAAAGT

ImpA104	[CA]_8_	F: TCTTTCTCTCTCTCCCACAC	53	60	MK742833

R: CTTCCGATCTGTAAAGAGA

ImpA105	T_9_	F: TGCAGAACTAATCAACATAGAGT	54	69	MK742834

R: CACAAAACGTTATGAATCTTTC

ImpA106	A_8_	F: TCAACAACTGAGCAAACCTAGA	57	60	MK742835

R: GTTGTTCATCTGTCCTTGGGA

ImpA107	A_9_	F: TTTCTTGAACTGAAATCCCAAA	55	53	MK742836

R: AAACTTCAACAACGTCTAACCAGA

ImpA108	T_8_	F: AATGGGGCAGTTAGTTTGGA	54	53	MK742837

R: AAGCAAATTTCCGAACAATT

ImpA109	T_10_	F: TGCAGTTCGAACAAGAAAATCTG	56	57	MK742838

R: CAGATTCTGGATTCATAATGATTGA

ImpA110	T_9_	F: CAGTTGCATTCTGGGCTTTT	56	53	MK742839

R: AGTTGGATCATCTCTCACGT

ImpA111	A_8_	F: TGCAGTTTTCCATTTTGTTTTG	52	69	MK742840

R: CCAAACAAAGTCATTCATTT

ImpA115	T_8_	F: GCAGAATCAAGTGGGGAAGG	58	68	MK742841

R: GCTCTTCCGATCTGTAATTTCCA

ImpA119	A_8_	F: TGCAGGATATGATGGCTCGC	58	69	MK742842

R: GCTCTTCCGATCTGTAACCT

ImpA120	A_12_	F: TGCAGCAAGATTTTCAAACT	54	56	MK742843

R: TCTTTACTCTTAGACGGAATGT

ImpA121	A_11_	F: TGCAGCAAGATTTTCAAACT	55	60	MK742844

R: TCCAGCCTTTACTTGTAAACAGA

ImpA122	[GA]_5_	F: TGAGAAACTATGTTGGGTCATGT	58	62	MK742845

R: TTCATGCTGCTGTCTCTCCT

ImpA127	A_8_	F: TCATCTCTGTGGCAAATTTCCA	55	56	MK742847

R: TCCGATCTGTAAATTCTTTATCA

ImpA128	T_8_	F: TGCAGTTGTTTTCTTGTGATTAGC	59	64	MK742848

R: CGATCTGTAACAACACTGCCC

ImpA130	T_8_	F: TGCAGAAAATGATATCAAGCTCT	56	58	MK742849

R: TGACATGATTCTTACTTAGCATAGG

ImpA131	[G/T]_19_	F: TGCAGACCACGATGTTCCT	56	69	MK742850

R: CAAACATCTTTTGCATAAACAAC

ImpA132	T_8_	F: TGCAGTATCGATGTTGATTGTGT	57	58	MK742851

R: ACAATTAGGTATCGTTGCAACT

ImpA133	A_8_	F: ACTTGTGAGCTATTCGATCGG	56	61	MK742852

R: TCTGTAAACGAGCTTGAACA

ImpA134	A_8_	F: ACTTGTGAGCTATTCGATCGG	58	61	MK742853

R: TCTGTAAATGAGCTTGAACGAGT

ImpA135	T_9_	F: GCTGTGATCTTCAATGGATGT	54	57	MK742854

R: TGTATATCTAGCTGTTATCCAA

ImpA137	A_8_	F: AGAGAAGAAATGATGCATGAGGT 56	66	MK742855

R: TCTTCCGATCTGTAATGTTTTCT

ImpA139	T_9_	F: TGCAGAAGACTTAGTATGTTTCTC 57	69	MK742856

R: ACCGCTCTTCCGATCTGTAA

ImpA141	A_10_	F: TGCAGCAAGATTTTCAAACT	54	54	MK742857

R: TCTTTACTTGTAGACGGAATGT

ImpA144	A_8_	F: GGATCAACAGAAGATACACA	53	57	MK742858

R: TCATCTTCTTCATCCCCTTGT

ImpA146	[GAT]_11_	F: TGCAGTTCAATCATCAATATG	53	68	MK742859

R: CGTCGTCGTCATCATCAT

ImpA148	T_10_	F: TGGCTGTGATCTTCAATGGA	53	54	MK742861

R: TCTAGCTGTTATCCAAATAGAT

ImpA149	T_8_	F: CAACATTTGACAACCGCGT	55	61	MK742862

R: TGAAATGACGCTAAGTTGTT

ImpA150	T_8_	F: CAACATTTGACAATCGCGT	53	61	MK742863

R: TGAAATGACACTAAATCGTT

ImpA151	T_8_	F: ACATTTGACAACCGCGACTT	55	59	MK742864

R: TGAAATGACGCTAAGTTGTTT

ImpA152	T_11_	F: TGGCTGTGATCTTCAATGGA	53	55	MK742865

R: TCTAGCTGTTATCCAAATAGAT

ImpA153	T_11_	F: TGGCTGTGATTTTCAATGGA	53	55	MK742866

R: TCTAGCTGTTATCCAAATAGA

ImpA154	T_12_	F: TGGCTGTGATTTTCAATGGA	53	56	MK742867

R: TCTAGCTGTTATCCAAATAGA

ImpA155	T_12_	F: GTTGGCTGTGATCTTCAATGT	54	58	MK742868

R: TCTAGCTGTTATCCAAATAGAT

ImpA156	T_8_	F: TGGCTGTGATCTTCAATGGA	53	56	MK742869

R: TCTAGCTGTTATCCAAATAGAT

ImpB1	A_11_	F: TGCAGAAATGAAACAACAAT	53	56	MK756124

R: ACTACACTCACGGATAACCT

ImpB2	A_8_	F: TGCAGAAGGATTTATGAAGAA	54	60	MK756125

R: CGATACGAAATAAGTCAGTGTTGA

ImpB3	[GA]_6_	F: TGCAGACACTATTACAGAAG	52	56	MK756126

R: AATGGGTTATTCTCTTTCTT

ImpB4	A_8_	F: TGCAGACTAACTCTCAGAAGA	56	69	MK756127

R: AGGATGGACAATAGGCTTCTTCT

ImpB5	[GA]_5_	F: TGCAGAGAAGAGAAGTTGTAGGA 58	66	MK756128

R: CCTCCCCATAAACCCTCCTT

ImpB6	[TGG]_6_	F: TGCAGAGATGACCGCCGG	60	53	MK756129

R: ACTTCAAGCACTCTACCACCA

ImpB7	A_10_	F: GCAGAGCAAAGTAAAGAATT	53	68	MK756130

R: TGGGAGAGGAATGTTACTTT

ImpB8	T_10_	F: GCAGAGGTTTGAGATGATTGTGA	58	67	MK756131

R: GTGTTCTCATCGGCCTGAAA

ImpB9	T_8_	F: TTTTCAGCTAAACCTACGTT	54	53	MK756132

R: TTCAGCTAAAAGTCTCAGTCGA

ImpB10	T_8_	F: TGAAACAGAGAGAAAGCGTA	55	57	MK756133

R: GGGTCATGCTAGTTTCCTACA

ImpB11	[TA]_5_	F: AGCTGACTGTTCTTTCTATCCT	57	61	MK756134

R: CGGCCTTACACTAAAATACCTGG

ImpB12	[A/G]_25_	F:_:_ CAGTGGCGATCGAGAGGAG	59	55	MK756135

R: CGAAACCCTACTTTGATCACTCTC

ImpB13	[GAG]_5_	F: GCATATGGACAACAACAGGGA	55	54	MK756136

R: AGACCAATTATTATTACTACCTCC

ImpB14	A_8_	F: TTCCCCATTTGGTGCATGC	56	63	MK756137

R: GGTTTTGTCTAGATTTTGGCT

ImpB15	T_8_	F: TGCAGGAAAGATGTTGATTT	54	65	MK756138

R: CAGTAAGAAAAGAAAGTCAACAACA

ImpB16	A_8_	F: AAGTTGGCCACACTCAATCG	58	62	MK756139

R: CGATCTGTAATCCTGAATTTCATCA

ImpB17	T_9_	F: TGCAGGTTAGTAGTATTCTTT	52	63	MK756140

R: TGTAAACAGTCACAATAACATGA

ImpB18	A_8_	F: TTTCTTGAACTGAAATCCCA	54	53	MK756141

R: GAAACTTCAACAACGTCTAACCA

ImpB19	T_9_	F: TGCAGGTTTGGCTTCTGG	58	67	MK756142

R: ACCGAACAGCTTGATTCATTCT

ImpB20	[CATA]_5_	F: TGCAGTAATTTGATAGCAA	52	57	MK756143

R: TCTGCTCATCTTCTAGGGTA

ImpB21	[CAT]_5_	F: GCAGTAGTAGCAACAGTGGA	57	66	MK756144

R: TCATCTTCTTCCTCCTCGCC

ImpB27	A_9_	F: TGCAGATCTTGTTGAAGGTT	53	69	MK756145

R: TCTTCCGATCTGTAAATGAT

ImpB30	[TAA]_5_	F: TGCAGCAAAGATAATACTAT	51	69	MK756146

R: TCCGATCTGTAAAGTATGTT

ImpB31	[C/T]_29_	F: ACCCTTCCCATTTCTCTCTC	53	57	MK756147

R: TCCGATCTGTAATTAGTAGAG

ImpB34	[AT]_7_	F: CCAATGGCCTTTCATGTCATCT	58	64	MK756148

R: CCGCTCTTCCGATCTGTAAA

ImpB35	A_8_	F: CCCATGCCAAGAATGTCACA	55	64	MK756149

R: TCCGATCTGTAAACACCATT

ImpB36	A_10_	F: GCGACTTTTCATCAACCCCA	54	65	MK756150

R: TCTTCCGATCTGTAAAACAATT

ImpB40	T_9_	F: TTGTTACGCAGCCAAAGTAA	53	63	MK756151

R: TCCGATCTGTAACTTTAGG

ImpB42	A_8_	F: TGCAGGATATGATGGCTC	54	69	MK756152

R: GCTCTTCCGATCTGTAATGT

ImpB45	T_9_	F: CTCCCGAGGTATAATCAC	52	60	MK756153

R: CGCTCTTCCGATCTGTAAAA

ImpB52	A_~~8~~_	F: TGCAGAATAAAGGAGGCATAA	56	53	MK756154

R: TGTCTCTAGAAATGAAGCCTGGT

ImpB54	T_9_	F: GCAGAGAGGCTATGTGATGT	58	67	MK756155

R: CGCTCTTCCGATCTGTAAAGC

ImpB56	[A/T]_31_	F: TGCAGATTTGACGGAAAA	54	67	MK756156

R: TCCGATCTGTAAATCTCAAAACT

ImpB57	[ATT]_6_	F: CCAGCATCTTCGAAACCAG	53	64	MK756157

R: CTCTTCCGATCTGTAAATAATAA

ImpB58	[A/T]_22_	F: TGCAGGTAATTTTCTCCCAT	54	69	MK756158

R: CTCTTCCGATCTGTAACACT

ImpB61	[GA]_9_	F. AGTTTGTGTGTGTTTGTGTGT	54	56	MK756159

R: ACGTATTTACTGTGTATTCTCT

ImpB62	[GAAA]_5_	F. TGCAGAAGATGAAGAAGAAAGA	56	65	MK756160

R: TGTTGTCAGATGAAGGGGAC

ImpB67	[TA]_6_	F: CAGCACAGAATAAGACAGGACA	57	63	MK756161

R: TCCAACTCCAAACTCCAATCA

ImpB68	[CAT]_6_	F: TGCAGTCCCAACCAACCG	59	69	MK756162

R: TGGAGAGAAGGAATGATGGTGA

ImpB69	[CTT]_5_	F: TGCAGTTCGTCTTCATTT	53	60	MK756163

R: TGTAGGGCTTATGAACAAGA

ImpB72	A_9_	F: TGCAGTCAGAAATAAGTATGAA	54	69	MK756164

R: CCGCTCTTCCGATCTGTAAC

ImpB73	[GA]_9_	F: TGCAGAAATGGTAGAGAGAG	55	56	MK756165

R: CCATTTTCATTTCTGCACAAACCT

ImpB74	T_10_	F: ATAACCGTCTTTAGTGAGCG	53	62	MK756166

R: GTTGATTGATTTGTAACAGAC

ImpB75	A_13_	F: TGCAGACCTGTTTGTTTT	53	59	MK756167

R: GGCATGGGATGGGAAGAC

ImpB76	[TA]_6_	F: TGCAGATGAATGATGAATGATGA	55	60	MK756168

R: TGCATGCACCAAAATATCTT

ImpB77	A_10_	F: TGCAGCAATCCAAAAGCATA	54	69	MK756169

R: CCGATCTGTAATTTCTCCTT

ImpB78	A_9_	F: TGTAGAATGATTGACTAGCCT	53	55	MK756170

R: CCGATCTGTAAAACTAAATGT

ImpB79	T_9_	F: GTCGCCTGGAAGCTTGATAC	57	60	MK756171

R: TCAAATTCCATTTACCAACTCCA

ImpB80	[CAAAA]_5_	F: TGCAGCTCATTGAAACATAACA	54	68	MK756172

R: TCTTCTTCTGTATTTTCTCTGT

ImpB81	A_10_	F: CTGCCTGAAACCCGATCAAA	56	64	MK756173

R: TCTTCTTAGTCTTCAAGTTCAGT

ImpB82	A_8_	F: GCAGCTGGATCAACGGAAG	58	67	MK756174

R: TCATCTTCTTCTTCATCCTCTTGA

ImpB83	A_9_	F: TGCCAAATTCTTCATTCTGC	53	56	MK756175

R: GAAGATATTCCTAATACAACATC

ImpB84	A_8_	F: TGCCAAATTCTTCATTCTGC	53	56	MK756176

R: GAAGATATTCCTAATACAACATC

ImpB85	A_9_	F: CACCTTTGGAGCGTCTAAA	53	61	MK756177

R: AGGAAAACTTGATACACATT

ImpB86	T_12_	F: TGCAGAAAAGGACATGTTTT	55	69	MK756178

R: CGATCTGTAATTCCCTGTCACC

ImpB87	[T/A]_20_	F: CAGATGCTCTCGGTTAGA	52	64	MK756179

R: ATGGATGATCAAACACAAAA

ImpB88	T_9_	F: TGCAGCCCTACTATTTGAA	53	65	MK756180

R: CTGTAAGTTGTAGGTCCAA

ImpB89	[GA]_6_	F: CAGTGCGTGCTTGTGTGT	57	56	MK756181

R: TCTTCTTCATCTTCTCTCTCTATCT

ImpB90	T_10_	F: TCACTCGACCATTCAAGATT	54	58	MK756182

R: ACTGGTATCGATTCAAAGAAGA

ImpB91	[GA]_6_	F: GCAGAGAAGAGAAGTTGCAGG	58	58	MK756183

R: CAAGTCCAAGTCACAGTCACA

ImpB92	A_12_	F: CCATCAGGTTTGTTACAAAA	52	59	MK756184

R: CCGATCTGTAACTCTAGCTA

ImpB93	[TTC]_5_	F: TAGCTTCTCGTCGTCTTCTT	53	55	MK756185

R: CGATCTGTAAATTGAAAAGAAG

ImpB94	[CAT]_8_	F: TGCAGCAAGAAGATCCATCA	57	69	MK756186

R: AGGTAAGTCTCTTTCATACTGCT

ImpB95	[GA]_9_	F: TGCAGCTGACAATGAGAGAGA	57	60	MK756187

R: TCTCTTCTCTATATTGGCGTCT

ImpB96	[CA]_11_	F: TGCAGAAAACTGAATGAATA	51	69	MK756187

R: TTCCGATCTGTAACCTTTT

ImpB97	A_11_	F: TGCAGAAAACTGAATGAATA	51	69	MK756189

R: TTCCGATCTGTAACCTTTT

ImpB111	A_9_	F: TGCAGGATATGATGGCTC	53	69	MK784667

R: CTCTTCCGATCTGTAATGTT

ImpB112	T_10_	F: CTCCCGAGGTATAATCAC	52	61	MK784668

R: CGCTCTTCCGATCTGTAAAA

ImpB113	A_8_	F: GCAGTCATGTCATCTAATCAGGA	55	68	MK784669

R: CTTCCGATCTGTAAACACAA

ImpB120	T_9_	F: CCAAGTGGGGAAGGAAAATGT	56	62	MK784670

R: CTCTTCCGATCTGTAATTTCCA

ImpB121	A_11_	F: TGCAGTAAACGTTGTCAAAC	56	69	MK784671

R: CGCTCTTCCGATCTGTAAAAG

ImpB123	T_10_	F: TGCAGAAACATTCAAAGCCT	56	60	MK784672

R: GGCATCTCTCAAGGTTGTGT

ImpB124	[T/A]_23_	F: ACTGGTTGGTTCATTTAGGT	53	63	MK784673

R: AAATTACAACACTAAAGGCT

ImpB125	A_8_	F: CACTTCTGAAGGCCCATAAGC	56	58	MK784674

R: TCTGTAACTCATAGTAAAGTGATGA

ImpB126	A_10_	F: TGCAGATTACGACGTTATTT	54	65	MK784675

R: CGGTCATCGTCGTAATCTTT

ImpB127	A_14_	F: GCAGCCAACTAGAATGTCT	52	66	MK784676

R: TGTTTATTTTGACTGAGACA

ImpB128	T_8_	F: TGCAGCTCCTTGATCTCAA	55	66	MK784677

R: TGTCATCATTTGCATTTTAGAAG

ImpB129	[CT]_5_	F. TGCAGCTTTATTTTCTGATTCGT	54	60	MK784678

R. TTGTAATCAATGGAGTCAATTC

ImpB130	[CAT]_5_	F: GCAGGCGGAGGAGATCAG	58	65	MK784679

R: ACATTTCATTCTCTTTGTGATTGCT

ImpB131	A_8_	F: GGAAACCTGTCAACAAGATCA	53	54	MK784680

R: TCTTTGTTTTCGTGTTTGAA

ImpB132	T_11_	F: TGCAGTTAGTCCACCTTTT	55	66	MK784681

R: TGTGAGCAGTAGAGATATCAAACA

ImpB133	T_9_	F: AACTTGACCCCAAAGTAGAA	52	58	MK784682

R: TGTTCTCACTAAATCTGAAA

ImpB134	A_10_	F: TGCAGACCTGTTTGTTTGTGT	59	68	MK784683

R: ATGGCATGGGATGGGAAGAC

ImpB135	T_8_	F: TGAAACAGAGAGAAAGCGTA	52	57	MK784684

R: CATGCTAGTTTCCTACAATAA

ImpB136	[CTT]_7_	F: CAGAACTATGGCTGGCTAG	55	61	MK784685

R: AGTCTGGCCGAAGAAGAAGA

ImpB137	T_10_	F: TGCAGTCTGTCGGTTTATAAGA	57	68	MK784686

R: CCCCATTTGTCATCCCAAGT

ImpB138	[TA]_6_	F: TGCGCTGAATCCTTCTAAAG	53	62	MK784687

R: GATCTGTAATCGAAATGGTT

ImpB139	T_9_	F: TGCAGAGAATTTCAGGTATACT	53	69	MK784688

R: CTTCCGATCTGTAATGAATT

ImpB143	A_8_	F: TGCAGCTCCAACATGTTTCA	56	69	MK784689

R: TCCGATCTGTAACGACTGT

ImpB145	A_8_	F: GTTTGTCAACTCATAAGAAA	52	57	MK784690

R: CCGATCTGTAATATAATGAGCC

ImpB147	T_9_	F: TGCAGTAGTATAAGACAGTAA	52	69	MK784691

R: GCTCTTCCGATCTGTAAG

ImpB148	A_9_	F: TGCAGTTCTAGAAACTTTAGA	53	69	MK784692

R: TCTTCCGATCTGTAATGAAAAGT

ImpB149	T_9_	F: TGCAGAACTAATCAACATAGAGT	53	69	MK784693

R: CACAAAACGTTATGAATCTTTC

ImpB150	[TA]_5_	F: GCAGGGTTGGTAAGCTAGCT	57	68	MK784694

R: TTCATGTTCAAATACTCTCTCCT

ImpB151	A_9_	F: TTTCTTGAACTGAAATCCCAAA	55	53	MK784695

R: AAACTTCAACAACGTCTAACCAGA

ImpB152	A_8_	F: TGCAGTTTTCCATTTTGTTTTG	52	69	MK784696

R: CCAAACAAAGTCATTCATTT

ImpB155	T_8_	F. GCAGAATCAAGTGGGGAAGG	58	68	MK784697

R: GCTCTTCCGATCTGTAATTTCCA

ImpB156	[CAA]_6_	F: GCAGCAACAGCATTACAACA	57	68	MK784698

R: CCGCTCTTCCGATCTGTAAG

ImpB158	T_12_	F: CGAGCCAGCCAAACTCTATT	58	56	MK784699

R: CCGCTCTTCCGATCTGTAAA

ImpB159	T_11_	F: TGCAGTAATTCGTCCTAT	51	58	MK784700

R: TGTAATATCGCTTGCAGAA

ImpB160	[GAA]_8_	F: AGCTGAACTCCGATGAAGAAGA	58	63	MK784701

R: CGCTCTTCCGATCTGTAATCT

ImpB161	A_11_	F: TGCAGCAAGATTTTCAAACT	54	55	MK784702

R: TCTTTACTCTTAGACGGAATGT

ImpB162	A_9_	F: TGCAGCAAGATTTTCAAACT	54	53	MK784703

R: TCTTTACTTGTAGACGGAATGT

ImpB163	[GA]_5_	F: TGAGAAACTATGTTGGATCATGT	56	62	MK784704

R: TTCATGCTGCTGTCTCTCCT

ImpB165	T_8_	F: TGCAGTTGTTTTCTTGTGATTAGC	59	64	MK784705

R: CGATCTGTAACAACACTGCCC

ImpB166	T_8_	F: TGCAGAAACCACAACTCGAT	55	69	MK784706

R: TCTGTAATTTAGGTTGGTGCT

ImpB167	[TC]_5_	F: CCGTTGTCATTATTGCCTCTTCT	58	54	MK784707

R: AGAGAAACAGAGACTTGAGGAAAA

ImpB168	A_8_	F: ACTTGTGAGCTATTCGATCGG	56	61	MK784708

R: TCTGTAAACGAGCTTGAACA

ImpB170	A_8_	F: AGAGAAGAAATGATGCATGAGGT 56	66	MK784709

R: TCTTCCGATCTGTAATGTTTTCT

ImpB171	A_10_	F: TGCAGCAAGATTTTCAAACT	54	54	MK784710

R: TCTTTACTTGTAGACGGAATGT

ImpB172	[GAT]_11_	F: TGCAGTTCAATCATCAATATG	53	68	MK784711

R: CGTCGTCGTCATCATCAT

ImpB173	T_11_	F: TGGCTGTGATTTTCAATGGA	53	55	MK784712

R: TCTAGCTGTTATCCAAATAGA

ImpB174	T_13_	F: AGTTGGCTGTGATCTTCAAT	53	64	MK784713

R: TGTATATCTAGCTGTTATCCAA

ImpB175	T_12_	F: TGGCTGTGATTTTCAATGGA	53	56	MK784714

R: TCTAGCTGTTATCCAAATAGA

ImpB176	T_8_	F: TGCAGGTAAATGGTATAATGTCTCT 56	62	MK784715

R: ACTTTCTAATGGAGGCAAACA

ImpB178	A_9_	F: GGTGGTCCAATCTGAGAAAA	52	61	MK784716

R: CTGTAAAGTTGTGACAACT

ImpB180	[CAT]_6_	F: GCCAAAGATAAATACATATGATCGA 55	60	MK784717

R: TCCAAGAAATAGGAAGATGATGA

ImpB185	[GAA]_5_	F: GCTGAACTCCGATGAAGATGA	57	62	MK784718

R: TCCGATCTGTAATCTTCACTTCT

ImpB187	A_10_	F: TGCAGAGTACAAGATTTAGAAGA	56	68	MK784719

R: AGTCAAAGGTAATCAGTAATGTGGA

ImpB188	[A/G]_27_	F:_:_ TGCAGAGGTTACAGTAGAGAGAG 58	63	MK784720

R: TCCGATCTGTAAGCTCTCTAGT

ImpB190	A_9_	F: TGCAGTTTTCCATTTTGTAGAGA	54	69	MK784721

R: GCTCTTCCGATCTGTAAAA

ImpB193	[TA]_5_	F: TGAATTCCATCAGTAAAACAAGT	55	59	MK784722

R: CGCTCTTCCGATCTGTAAG

ImpB195	T_8_	F: AACCGTCTTTAGTGAGCGTC	55	59	MK784723

R: TGTTGATTGATTTGTAACAGAC

ImpB196	A_8_	F: TGCAGGGCCCAGGGAAAG	60	53	MK784724

R: TTTTGTTTTGACTGGCATGTGTAAA

ImpB198	[GCC]_5_	F: CATGCCGTCACCACCAATAT	58	53	MK784725

R: AGAAGATGCACTCCGACGG

ImpB199	A_8_	F: TGCAGATTATCAACTTTAGT	51	61	MK784726

R: TGGTAGCCTTGTGATAATGT

ImpB200	[CTT]_8_	F: CAGCCAGCCTTCTTCTTCT	57	67	MK784727

R: GAGAAAACGGTTACCTGGGT

ImpB201	T_9_	F: TGCAGCCCTACTATTTGAAAGC	56	64	MK784728

R: CTGTAAGTTGTAGGTCCAAAAT

ImpB202	A_12_	F: TGCAGGAAAAGTAATGGATGACA	57	68	MK784729

R: TGAACAAATCCCTAATCAAGAAGA

ImpB203	[GAA]_6_	F: TGCAGGTCTTCTCTTTTCCCA	59	68	MK784730

R: TCTGTAATACCGAAGCAGCCT

ImpB204	T_8_	F: GGTTTTACCGTAGCCTTTT	54	53	MK784731

R: TTCCGTCGGTCAAGAACAG

ImpB208	T_9_	F: AGTTTCTCACTCGGTCTGAA	54	66	MK784732

R: GCTCTTCCGATCTGTAATAG

ImpB209	[GA]_7_	F: CGAGGGGAAAGAGAAAAGGA	57	57	MK784733

R: CGTTTCCTTTCAACCTTTCTCTC

ImpB210	[CAT]_7_	F: CAGCAGCAACATTATCATCA	55	65	MK784734

R: CCGATCTGTAATAAGCTAGAATTGA

ImpB213	[GA]_14_	F: TGCAGAGGTTACACGAGAGA	57	67	MK784735

R: AGCAACAATTTCAAGCTCTCT

ImpB214	[A/C]_16_	F: AGTGGTCCAATCTGAGAAAA	52	57	MK784736

R: AGTTGTGACAACTTGTTT

ImpB215	[CAT]_6_	F: TCAGCAACTTTTCATCATCATCA	57	57	MK784737

R: GCAGGTGCGTGTTTGAGTAT

ImpB216	[GA]_5_	F: TGCAGAGAAGAGAAGTTGTAGGA 58	68	MK784738

R: CGCACCATAAAACCCACCTT

ImpB219	[A/G]_19_	F: AGGAAGGAACCAATGGAGCA	58	66	MK784739

R: CGCTCTTCCGATCTGTAAGG

ImpB222	T_8_	F: ACAGGGGAGATTCAATGAAA	54	57	MK784740

R: TCTTCCGATCTGTAATCCCA

ImpB223	A_10_	F: GCAGATTTACATAGATTTACATAGC 56	68	MK784741

R: CGCTCTTCCGATCTGTAATGA

ImpB224	T_8_	F: CAACATTTGACAACCGCGT	54	62	MK784742

R: TCTTCCGATCTGTAATTTGTT

ImpB225	A_10_	F: TGCAGTAAACGTTGTCAAAC	54	69	MK784743

R. GCTCTTCCGATCTGTAAAAG

ImpB226	T_8_	F: TTTTACAGAGGAGATTCAAA	51	61	MK784744

R: CTCTTCCGATCTGTAATCCA

ImpB227	[GA]_5_	F: CGTGTGCGTGAGTGTGTG	57	55	MK784745

R: TCTTCTTCATCTTCTCTCTCTATCT

ImpB228	A_8_	F: TGCAGTCACTAATCCATGAGC	57	53	MK784746

R: ATGGCCTAAATTTTCTGCTAAGA

ImpB229	[GAT]_9_	F: TGCAGTTCAATCATCGAT	53	62	MK784747

R: CGTCGTCGTCATCATCAT

_______________________________________________________________________________________________
